# Clownfishes are a genetic model of exceptional longevity and reveal molecular convergence in the evolution of lifespan

**DOI:** 10.1101/380709

**Authors:** Arne Sahm, Pedro Almaida-Pagan, Martin Bens, Mirko Mutalipassi, Alejandro Lucas-Sanchez, Jorge de Costa Ruiz, Matthias Görlach, Alessandro Cellerino

## Abstract

Standard evolutionary theories of aging postulate that reduced extrinsic mortality leads to evolution of longevity. Clownfishes of the genus Amphiprion live in a symbiotic relationship with sea anemones that provide protection from predation. We performed a survey and identified at least two species with lifespan of over 20 years. Given their small size and ease of captive reproduction, clownfishes lend themselves as experimental models of exceptional longevity.

To identify genetic correlates of exceptional longevity, we sequenced the transcriptomes of *Amphiprion percula* and *A. clarkii* and performed a scan for positively-selected genes (PSGs). These were compared with PSGs detected in long-lived mole rats and short-lived killifishes revealing convergent evolution in processes such as mitochondrial biogenesis. Among individual genes, the Mitochondrial Transcription Termination Factor 1 (*MTERF1*), was positively-selected in all three clades, whereas the Glutathione S-Transferase Kappa 1 (*GSTK1*) was under positive selection in two independent clades. For the latter, homology modelling strongly suggested that positive selection targeted enzymatically important residues.

These results indicate that specific pathways were recruited in independent lineages evolving an exceptionally extended or shortened lifespan and point to mito-nuclear balance as a key factor.

## Introduction

The lifespan of vertebrate species spans two orders of magnitude from few months for annual killifish (1) to several centuries for the greenland shark (2). Understanding the genetic architecture underlying these differences is a major challenge but may deliver new insights into the mechanisms controlling evolution of lifespan and human longevity.

Next-generation sequencing technology has revolutionized evolutionary genomics as it allows to obtain genome-scale sequence information for large number of species. A particularly useful approach to identify the genetic architecture of evolutionary novelties is the analysis of positive selection. This approach requires the comparison of the sequence of protein-coding genes in related clades where one of the clades evolved the trait of interest, in this specific case exceptional lifespan. To date, several different mammalian taxa/clades where analysed with this approach with the purpose of identifying sequence changes associated to evolution of longevity: the elephant, the bowhead whale, bats and mole-rats (3–8). These analyses delivered interesting candidate genes and pathways that underwent accelerated molecular evolution in coincidence with evolution of exceptional lifespan. A major drawback of this approach is that these long-living mammals are difficult or impossible to be kept in captivity and manipulated experimentally. This creates the need for a long-lived vertebrate that is small in size, easily adaptable to captive life, can be bred in large numbers and therefore represents a convenient experimental model organism.

Standard evolutionary theories of aging predict that low extrinsic mortality conditions lead to the evolution of slow senescence and increased lifespan. Some examples that confirm these theories are the exceptional longevity of vertebrate species under low predation risk since they are chemically protected (9, 10), adapted to an arboreal life (11) or found in protected environments such as caves, respectively. On the other hand, annual fishes of the genus *Nothobranchius* provide an example of how increased extrinsic mortality conditions lead to the evolution of accelerated senescence and short lifespan (12–14). Analysis of positive selection in annual killifishes revealed a potential link between the evolution of genes governing mitochondrial biogenesis and the evolution of lifespan (15).

All clownfish species (genus *Amphiprion*) evolved a specific adaptation that allows them to live in symbiosis with sea anemones. Symbiosis evolved in the last common ancestor of clownfish and clownfish represent a monophyletic group in the Pomacentridae family (damselfishes) (16). In the Indo-Pacific Ocean, clownfishes are found in association with one or more sea anemone species and a large variation in host usage exists (17–19). Fish that feel threatened by predators immediately seek protection by the anemone’s tentacles; without that symbiosis, fishes are readily attacked and predated (20–22). Therefore, clownfish are protected from predation through reduction of extrinsic mortality owing to the presence of anemones (23). Hence, the overall mortality rate of clownfish is low as compared to other coral reef fishes or other tropical species of Pomacentridae of the same size (20, 23- 26).

All clownfish are born as males and develop, through protandrous hermaphroditism, into females: in a colony, only the dominant pair contributes to the reproduction of the colony (27). Other individuals of the colony are non-breeding males. Studies in the wild have shown that natural mortality of adult clownfishes can be very low: during the period 2011–2013, the average biannual mortality rate *per capita* varied, depending on the study site, between 0.18 and 0.49 for juveniles, 0.09 and 0.44 for males, and 0.19 and 0.55 for females (28). Predatory pressure differs in different stages of adulthood and is increased for non-breeding males (20).

These fishes are small in size (less than 10 cm for the smallest species) and the closely-related species *A. percula* and *A. ocellaris* are popular and hardy aquarium fishes, are bred in large numbers for the aquarium trade, and are subject to selective breeding to fix specific pigmentation patterns so that a number of different captive strains are available. For these reasons, clownfishes could become the first experimental model for long-living vertebrates.

In order to identify the genetic basis of adaptations linked to clownfishes’ exceptional lifespans, we performed a positive selection analysis. This analysis requires the identification of the closest related taxon that does not possess the trait of interest in order to exclude events of positive selection that predate the evolution of this trait (29).

Other species of damselfish evolved an inter-specific mutualistic relationship with branching corals (30, 31). In this case, corals are used by fishes as shelter that can provide protection from predators and a safe area to egg laying (32, 33). Among the family Pomacentridae, *Chromis viridis* shows an interesting relationship with a wide range of scleractinians (34, 35). Despite the presence of favourable microhabitat, *C. viridis* are predated by a wide range of generalist predator species. Hixon and Carr (36) suggested there is a clear relationship among transient and benthic predators and damselfish mortality: damselfish that search for protection in the shelter from transient predators are susceptible to attack by resident benthic predators and vice versa. In the presence of both groups of predators, mortality increases dramatically due to the lack of available refuge that expose *Chromis* to intense predation (36). Therefore, *Chromis viridis* represent a well-suited outgroup for our analysis because it shares with clownfishes several general traits linked to benthic life and symbiosis with corals but it is subject to much higher predation rates (Fig. 1).

**Fig. 1.**
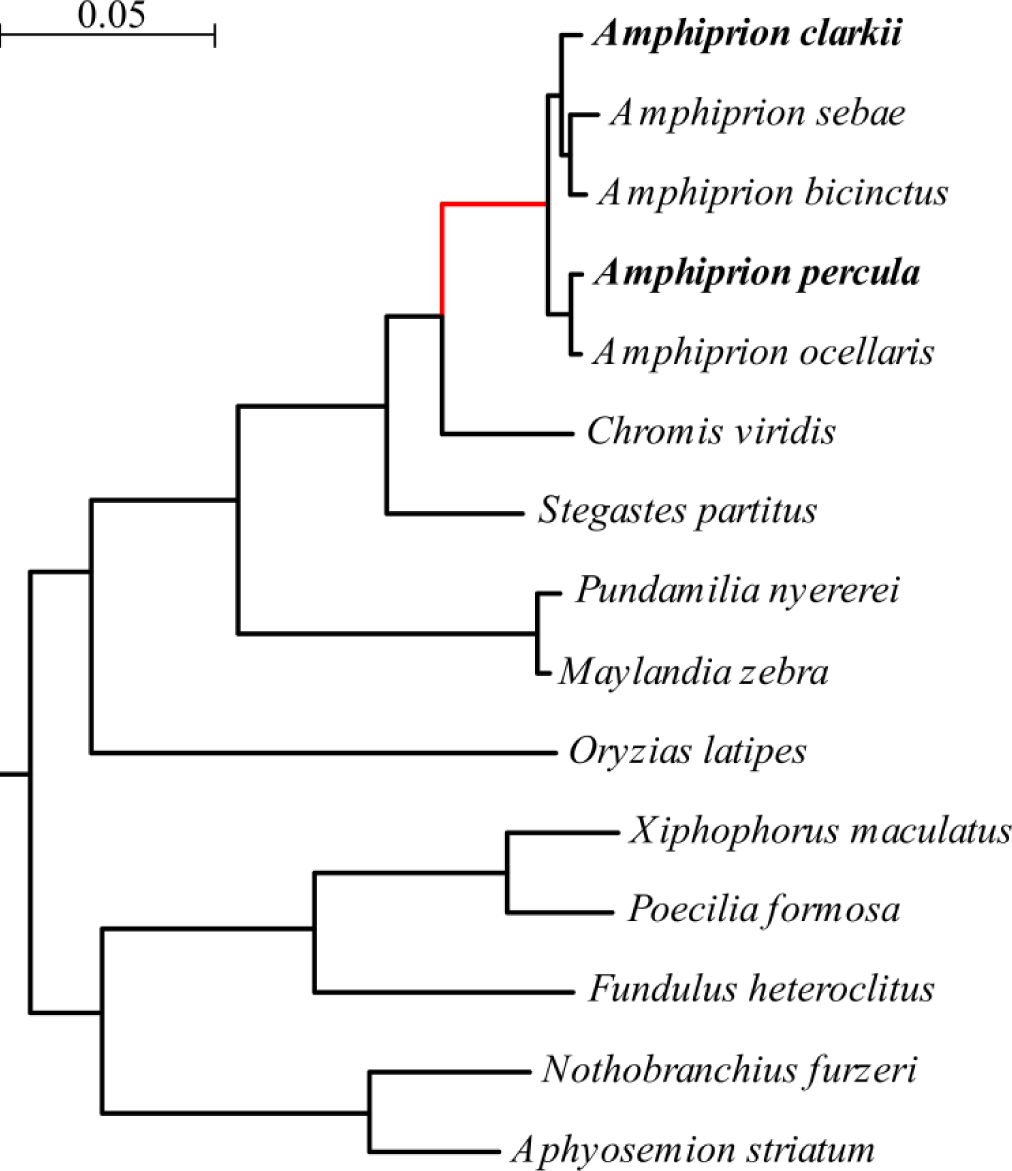
Nucleotide-based phylogeny of the analyzed fish species. We searched for positively selected genes on the last common ancestor of the clownfishes (*Amphiprion*, red). The two species *A. clarkii* and *A. percula* depicted in bold are those that were sequenced in this study. The phylogenetic tree was derived as part of the positive selection analysis with the PosiGene pipeline (62). Briefly, during this process 8215 genes were concatenated and the resulting concatenated alignment split in 404 fragments each of which had a length of 15 knt. From each fragment, a phylogeny was calculated via maximum likelihood and, from all resulting 404 trees, a consensus tree was determined using the Phylip package (72). The scale bar represents 0.05 substitutions per site.

## Captive lifespan of clownfishes

In order to obtain a reliable lower estimate for the captive lifespan of clownfish species, in 2016 we distributed a questionnaire to researchers working with clownfishes and to public aquaria across Europe (Table 1/S1), and surveyed existing literature. For six different species, at least one individual was reported to have lived more than 10 years and for two different species, *A. melanopus* and *A. ocellaris*, we obtained record of animals alive and actively spawning at an age of over 20 years. More systematic data could be obtained for the species *A. ocellaris* (the most common species in the aquarium trade). The oldest cohort for which a record was available comprised 27 fish born in 2008 of which 25 were still alive in 2016.

We conclude that there is solid evidence that at least the species *A. ocellaris* and *A. clarkii* can live in captivity for more than two decades making them the first teleost model of exceptional longevity.

**Table 1.** Results of the clownfish survey. The longest-lived individual for each species is indicated

| Species | oldest animal | status at census | size of group |
| --- | --- | --- | --- |
| *A. akydinos* | 13 | dead | 1 |
| *A. clarkii* (wild)* | 12 | alive | n.a. |
| *A. clarkii* (privately owned) | 16 | alive | 2/0 dead |
| *A. clarkii* | 9 | alive | 2/0 dead |
| *A. frenatus* ** | 18 | dead | n.a. |
| *A. melanopus* | 21 | alive | 2/0 dead |
| *A. ocellaris* (privately owned) | 22 | alive | 2/0 dead |
| *A. ocellaris* | 17 | alive | 2/0 dead |
| *A. perideraion* ** | 18 | alive | n.a. |

\* Moyer, 1986 (37)
\*\* Fautin and Allen, 1992 (17)

## Analysis of positive selection

In order to perform genome-wide scans for positive selection, we obtained the transcriptomes of the species *A. clarkii* and *A. percula* based on own sequencing using methods previously described for the killifishes (15). Furthermore, we assembled clownfish transcriptomes from public read data of *A. bicinctus*, *A. ocellaris* and *A. sebae*. As the closest-related non-symbiotic species, we additionally sequenced the transcriptome of *Chromis viridis*, a very abundant species in coral reefs. More distant outgroups were a selection of species from the series Ovalentaria, whose genomes are available in GenBank (see also (15)). We analysed positive selection on the branch leading to the last common ancestor (LCA) of all clownfish species (Fig. 1).

A total of 157 positively selected genes (PSGs) of 14214 analyzed genes were identified in the LCA of the clownfishes (Table S2). We tested for overrepresentation of gene ontology (GO, FDR <0.1) and observed 19 biological processes enriched for PSGs (Table 2, Table S3). A majority of these processes is of particular interest for aging research: altogether nine enriched processes are linked to the metabolism of xenobiotics, detoxification or glutathione metabolism, respectively. Interestingly, these processes were shown to be strongly up-regulated in experimental conditions favoring longevity such as dietary restriction and inhibition of the somatotropic axis making the animals more resistant to toxins (38–41). Furthermore, experimental manipulation of mitochondrial translation, another enriched process, is known to increase lifespan in *C. elegans* (42).

**Table 2.** Biological gene ontology processes enriched for positively selected genes (FDR<0.1).

| GOBPID* | Term | FDR** |
| --- | --- | --- |
| GO:1901685 | glutathione derivative metabolic process | 0.020646 |
| GO:1901687 | glutathione derivative biosynthetic process | 0.020646 |
| GO:0006805 | xenobiotic metabolic process | 0.08376 |
| GO:0032543 | mitochondrial translation | 0.08376 |
| GO:0071466 | cellular response to xenobiotic stimulus | 0.08376 |
| GO:0009410 | response to xenobiotic stimulus | 0.08376 |
| GO:0042178 | xenobiotic catabolic process | 0.08376 |
| GO:0007157 | heterophilic cell-cell adhesion via plasma membrane cell adhesion molecules | 0.08376 |
| GO:0050900 | leukocyte migration | 0.08376 |
| GO:0045321 | leukocyte activation | 0.08376 |
| GO:0007155 | cell adhesion | 0.08376 |
| GO:0022610 | biological adhesion | 0.08376 |
| GO:0048870 | cell motility | 0.08376 |
| GO:0051674 | localization of cell | 0.08376 |
| GO:1990748 | cellular detoxification | 0.08376 |
| GO:0055081 | anion homeostasis | 0.08376 |
| GO:0007229 | integrin-mediated signaling pathway | 0.085657 |
| GO:0016477 | cell migration | 0.085657 |
| GO:0098754 | detoxification | 0.085657 |

\* GOBPID – gene ontology biological process ID
\* FDR – false discovery rate (adjusted p-value for multiple testing)

Furthermore, mitochondrial translation was one of the mitochondrial biogenesis processes that were found to be enriched for PSGs in extremely short-lived killifishes (15). Recent observations of similar genes and pathways found to be affected by positive selection, both, in very long- and short-lived species led to hypotheses of antiparallel evolution (43, 44). This means that functionally opposite selection pressures with regard to the tradeoff between fast growth and a long lifespan can result in adaptations of the same genes and pathways – in opposite functional directions. We further tested this hypothesis by using Fisher’s method to combine enrichment p-values across the results of the recent positive selection analyses in short-lived killifishes and the analysis in clownfishes. In this meta-analysis, 34 genes exhibited a signature of positive selection (FDR<0.1) across species (Tables S4-S6).

**Table 3.**
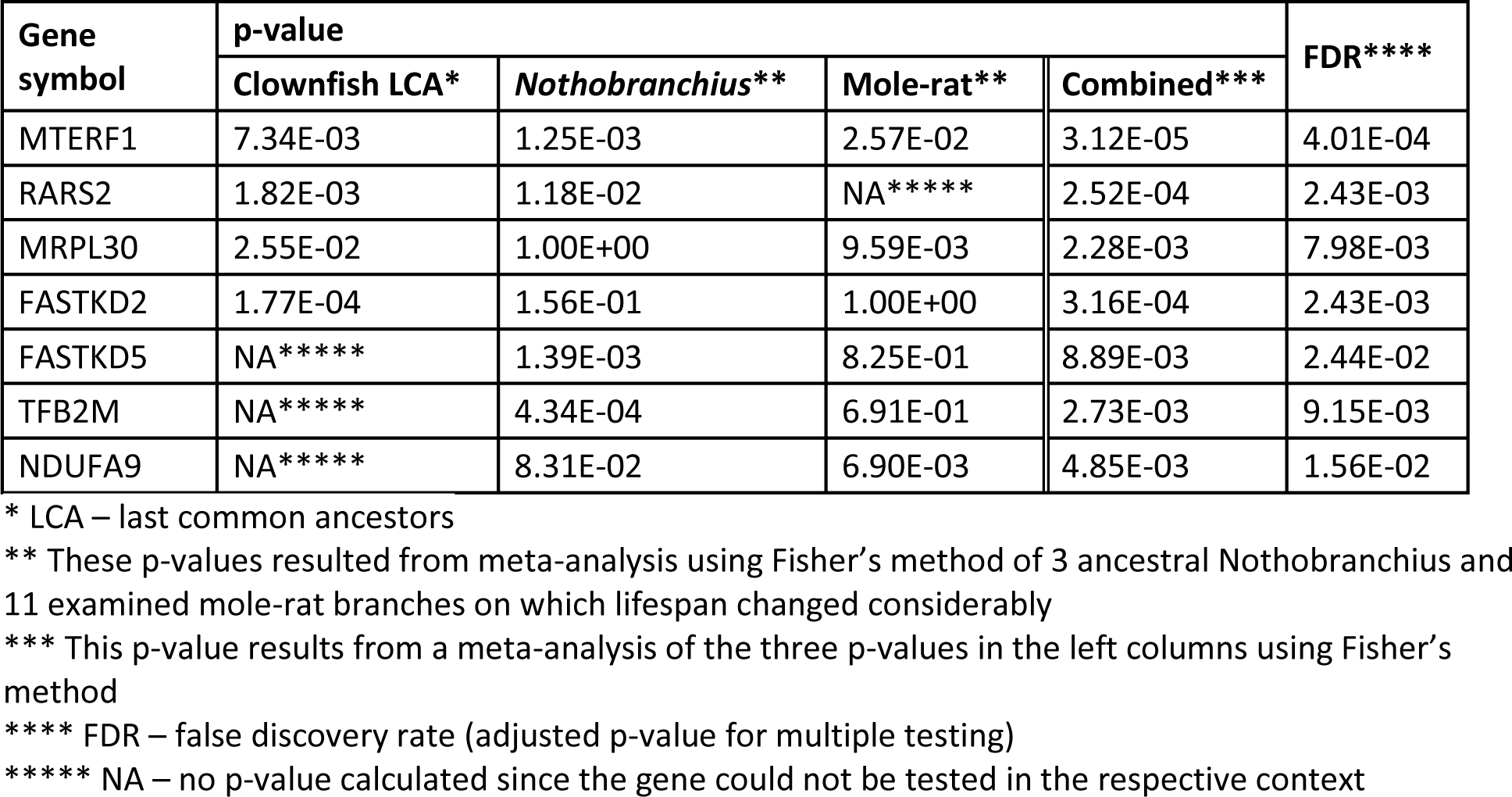
Positively selected genes associated with mitochondrial biogenesis identified in a meta-analysis across three evolutionary clades with exceptional short or long lifespans.

| Gene<br>symbol | p-value |  |  |  | FDR**** |
| --- | --- | --- | --- | --- | --- |
|  | Clownfish LCA* | *Nothobranchius* ** | Mole-rat** | Combined*** |  |
| MTERF1 | 7.34E-03 | 1.25E-03 | 2.57E-02 | 3.12E-05 | 4.01E-04 |
| RARS2 | 1.82E-03 | 1.18E-02 | NA***** | 2.52E-04 | 2.43E-03 |
| MRPL30 | 2.55E-02 | 1.00E+00 | 9.59E-03 | 2.28E-03 | 7.98E-03 |
| FASTKD2 | 1.77E-04 | 1.56E-01 | 1.00E+00 | 3.16E-04 | 2.43E-03 |
| FASTKD5 | NA***** | 1.39E-03 | 8.25E-01 | 8.89E-03 | 2.44E-02 |
| TFB2M | NA***** | 4.34E-04 | 6.91E-01 | 2.73E-03 | 9.15E-03 |
| NDUFA9 | NA***** | 8.31E-02 | 6.90E-03 | 4.85E-03 | 1.56E-02 |

\* LCA – last common ancestors
\*\* These p-values resulted from meta-analysis using Fisher's method of 3 ancestral *Nothobranchius* and 11 examined mole-rat branches on which lifespan changed considerably
\*\*\* This p-value results from a meta-analysis of the three p-values in the left columns using Fisher's method
\*\*\*\* FDR – false discovery rate (adjusted p-value for multiple testing)
\*\*\*\*\* NA – no p-value calculated since the gene could not be tested in the respective context

An overrepresentation analysis of GO terms among the genes yielded signatures of positive selection in the meta-analysis across different species with exceptional lifespans. As in our previous examination of short-lived killifishes (15), we found an enrichment for mitochondrial biogenesis functions (p=1.05*10^−5^, Table S7). Among the genes involved in mitochondrial biogenesis were *TFB2M* and *MTERF*, that are necessary for mitochondrial transcription, *FASTKD5* and *FASTKD2* whose gene products are required for the biogenesis of mitochondrial ribosomes, (45), as well as *RARS2* coding for a mitochondrial tRNA-synthetase.

Among the other 15 PSGs genes showing evidence for positive selection, both, in the clownfish LCA and in meta-analysis were, e.g., *LAMP2* and *CD63* (also called *LAMP3*) which code for major protein components of the lysosomal membrane (46, 47). In addition, *CD63* gene expression was shown to predict the malignancy grade of many different tumor types (48–52) and the artificial prevention of the decrease of *LAMP2* gene expression during aging in mice results in considerably reduced cell damage, as well as in liver functions in old mice that are indistinguishable from those in young mice (53). Finally, another interesting example that was identified as significant, both, in the clownfish LCA and in the meta-analysis, is *GSTK1* encoding a glutathione-S-transferase that localizes to the peroxisome. GSTK1 was shown to be associated with diabetes type 2 which is another major aging related disease (54, 55).

The positive selection analysis provides not only candidate genes but also candidate amino acids for follow-up studies. To exemplify this, we performed protein homology modeling for GSTK1 starting from the publicly available structures of the human dimeric apoenzyme (PDB 3RPP, (56)) and the rat dimeric enzyme with the bound GSH substrate (PDB 1R4W; (57)). The latter was used to assess on a structural basis the relationship of the six positively selected sites in the clownfish with those that are known to be involved in the enzyme’s function (57). Interestingly, also the LCA of Nothobranchius shows positive selection in GSTK1 contains, in addition, one site with high probability of positive selection in the LCA of Nothobranchius (Glu167, blue in Fig.2). The selected site in Nothobranchius, however, is structurally remote to the functionally relevant sites. In contrast, we found that in clownfish two of three sites that were predicted with high probability to be positively selected (≥ 95%, Phe60, Met63, red in Fig.2) and one of three sites with lower probability (41%, His64, orange in Fig.2) belong to the same α-helical stretch of amino acids that lines the substrate access channel, contribute to the dimer interface (Asn61, Tyr65, Asp69, green in Fig. 2) as well as to the substrate binding sites (Lys62, turquoise in Fig. 2), respectively (57). The third site with a high probability to be positively selected is Glu88 (brown in Fig 2). Glu88 is one of four amino acids at the entrance of the substrate access channel and situated in close proximity to Pro55, Pro56 and Pro87 (black in Fig. 2). The latter three are also part of the substrate access channel (57). We found another site positively selected with a lower probability in close proximity to the dimer interface (Lys177, orange in Fig. 2). This positive selection at particular positions related to enzymatic function invites the speculation that it might have a bearing on the enzymatic activity of the clownfish GSTK1, but this hypothesis would have to be tested experimentally.

**Fig. 2.**
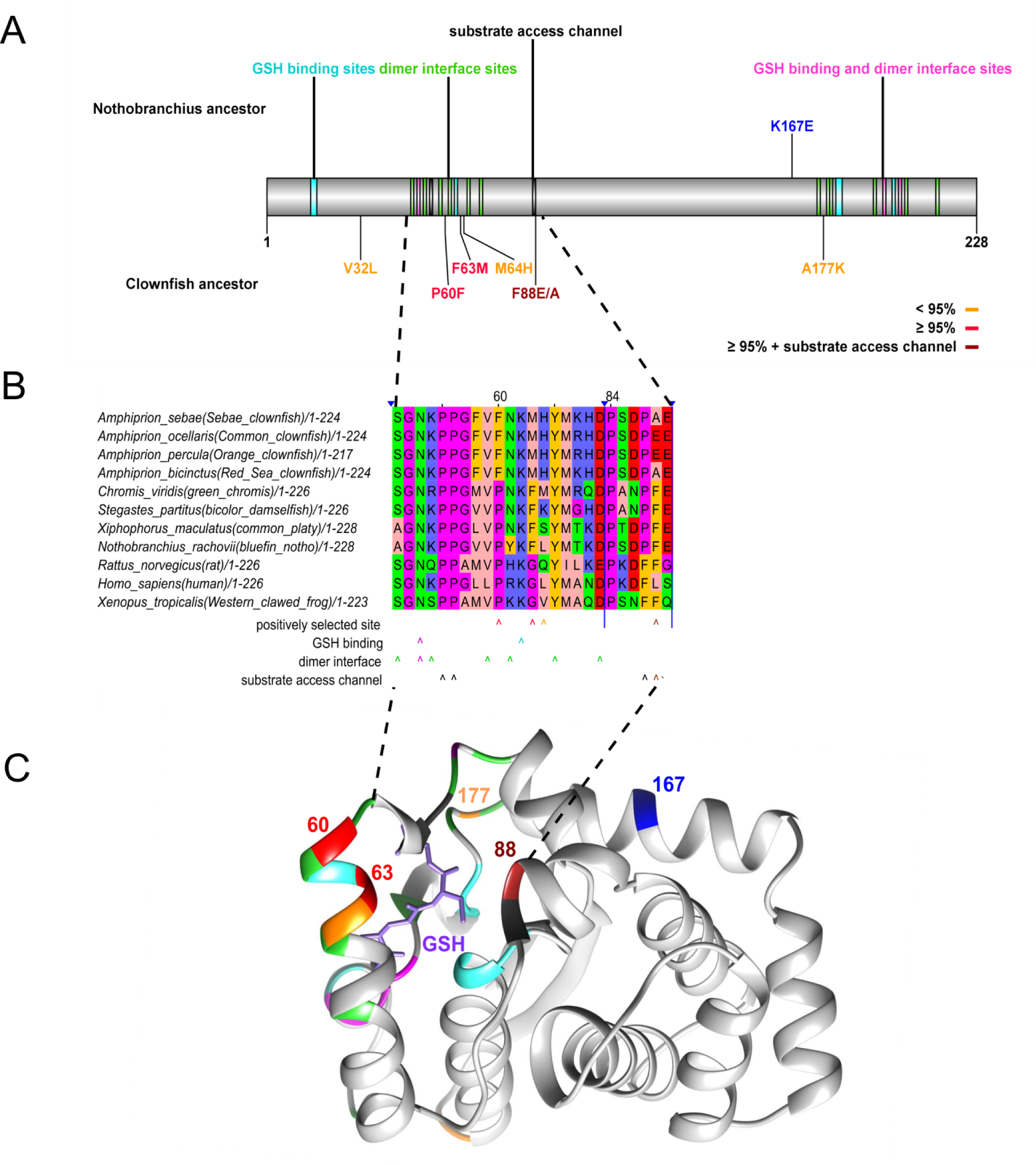
(**A**) Linear depiction of GSTK1 with color coded known functional domains/sites (dimer interface – green, GSH binding – turquoise, sites that serve as both dimer interface and GSH binding – violet, substrate access channel – black) and positively selected sites (in the last common ancestor of the clownfishes with a predicted probability ≥ 95% – red, in the last common ancestor of the clownfishes with a predicted probability < 95% – orange, in the last common ancestor of *Nothobranchius pienaari* and *Nothobranchius rachovii* – blue. (**B**) Alignment of GSTK1 orthologs across a wide phylogenetic range of species. Depicted are two protein regions (51-69, 84-89) that contain positively selected sites and functionally relevant sites in close proximity. The color code for positively selected and functionally relevant sites is the same as in panel A. (**C**) Clownfish GSTK1 model showing one subunit of the modelled dimer (for an overview see SI Fig S1. Selected positions are color coded according function depicted in the overview scheme at the top. The numbered and colored residue positions (60, 68, 88, 170 and 177) are discussed in detail in the text. Also shown is the GSH substrate (glutathione, light purple) as positioned in the template structure (PDB 1R4W) of the rat GSTK1.

## Conclusions

We have provided evidence for exceptional longevity of clownfishes in captivity. The species *A. ocellaris* is bred in captivity and commercially available in large numbers and we suggest this species as laboratory model for extended lifespan.

Analysis of positive selection has shown evolutionary convergence both with the exceptionally short-lived genus *Nothobranchius* and with exceptionally-long lived mole rats.

In particular, clownfishes and mole rats both show positive selection in two key proteins of the lysosome: LAMP2 and CD63. These results are consistent with the conserved up-regulation across tissues and species of genes coding for lysosomal proteins and widespread accumulation of lysosomal aggregates observed during aging (58, 59) and suggests that lysosomal function is of key importance for evolution of exceptional longevity. Another interesting example of convergent evolution is GSTK1, which is positively selected in both the exceptionally-long and exceptionally-short lived fish clades. GSTK1 is involved in glutathione metabolism. Since detailed structures of this protein are available (56, 57), homology modelling was possible and it strongly suggests that positive selection targeted positions that are involved in the enzymatic function of the encoded protein.

Finally, prominent signs of convergence were observed for genes and pathways related to biogenesis of mitochondrially-encoded proteins with the remarkable observation that MTERF is under positive selection in all three taxa. These findings point to the key importance of mito-nuclear balance in the regulation of animal longevity.

## Methods

### Clownfish lifespan estimation

The determination of clownfish lifespan was performed through the distribution of an internet-based questionnaire to zoos and aquariums worldwide, requesting information on clownfish demographic details: (1) the various clownfish species maintained in captivity, (2) the number of individuals for each species, (3) if each individual is captive bred or not, (4) the year of acquisition and, if not still alive, death, and (5) the sex of each individual, if determined. The questionnaire was circulated in 2016 to international associations and organizations of zoos and public aquariums such as the European Association of Zoos and Aquaria (EAZA), the Association of Zoos and Aquariums (AZA), the European Union of Aquarium Curators (EUAC) and the World Association of Zoos and Aquariums (WAZA).

Responses to our questionnaire were received from 5 zoos and aquariums as well as two private entities (see Acknowledgments).

### Experimental fish and sampling

Sub adult *Amphiprion percula* (total length, 45.2±1.2 mm; Wt, 1.6±0.1 g, n=12), *Amphiprion clarkii* (total length, 46.4±5.1 mm; Wt, 2.3±0.9 g, n=12) and *Chromis viridis* (total length, 43.0±1.6 mm; Wt, 1.3±0.1 g, n=12), were used. Animals were acquired from local dealers and subjected to acclimation during one month in the facilities of the Marine Acquarium at the University of Murcia (Spain). Fish were kept in groups under exactly the same conditions (temperature, 27±1 ºC; salinity, 24±1, pH, 8±0.2; dissolved oxygen, 6.5±0.2 mg/L) and fed *ad libitum* four times a day a standard low-fat diet to match their requirements (composed by Mysis shrimp, enriched *Artemia nauplii* and red plankton).

Fish were euthanized by exposure to the anesthetic benzocaine hydrochloride (400 mg l-1) for 10 min following the cessation of opercular movement. Brains, livers and samples of skeletal muscle were collected for analyses. For each species, three whole brains were frozen in l-N and stored at −80 ºC prior to molecular determinations.

The animal procedures were approved by responsible authorities (A13160603, from the Consejeria de Agua, Agricultura, Ganaderia y Pesca, Comunidad Autonoma de la Region de Murcia, Spain).

### Coding sequence data

Our analysis comprised five clownfish species (*A. ocellaris*, *A. clarkii*, *A. bicinctus*, *A. percula*, *A. sebae*), *C. viridis* representing the non-symbiotic sister-taxon of the *Amphiprion* genus and nine more distantly related outgroup species (*Stegastes partitus*, *Pundamilia nyererei*, *Maylandia zebra*, *Oryzias latipes*, *Xiphophorus maculatus*, *Poecilia formosa*, *Fundulus heteroclitus*, *Nothobranchius furzeri*, *Aphyosemion striatum*). mRNA sequences of the ougroups were obtained obtained from RefSeq along with their coding sequence annotation (Table S8). For *A. ocellaris*, *A. bicinctus*, *A. sebae* we downloaded read data from the short read archive (Bio projects PRJNA374650, PRJNA261388 and PRJNA285007, respectively). For *A. clarkii*, *A. percula* and *C. viridis* we performed novel RNA-seq as described in Table S9. The reads of the clownfishes and *C. viridis* were preprocessed using SeqPrep with minimum adapter length of five as well as a demanded minimum read length of 50. *De novo* transcriptome assemblies for these species were performed using FRAMA with *Stegastes partitus* as reference species (60). For the clownfishes and *C. viridis* the longest isoform was chosen to represent the gene. For the outgroups, in cases in which multiple isoforms per gene were annotated based on the reference, all of them were used in subsequent analyses. The assembly completeness of all examined species were estimated using BUSCO (61), was 90-100% (Table S8).

### Identification of positively selected genes

To scan on a genome-wide scale for genes under positive selection, we fed the coding sequences of the described species set into the PosiGene pipeline (62). *Stegastes partitus* was used as PosiGene’s anchor species. Orthology was determined by PosiGene via best bidirectional BLAST searches (63, 64) against *Stegastes partitus*. The branch of the last common ancestor of the clownfishes was tested for genes under positive selection (Table S2). FDR <0.05 was used as threshold for significance.

### Gene ontologies

We determined enrichments for GO categories using Fisher’s exact test based on the R package GOstats (Table S3). The resulting p-values were corrected using the Benjamini-Hochberg method (65). We used throughout the manuscript 0.1 as significance threshold. Enrichment for mitochondrial biogenesis genes was tested using Fisher’s exact test and the union set of the genes in the following five mitochondrial related GO terms: GO:0000959, 0032543, 0045333, 0033108, 0070584 (Table S5). The same GO terms were used in our previous study (15) to test for enrichment

### Meta analysis

To identify genes that show signs of positive selection across multiple evolutionary branches on which lifespan was altered considerably, we combined p-values from this study with those of two previous studies using Fisher’s method (66) (Table S4-S6). In all three studies, PosiGene was used to determine p-values. The first study searched for genes under positive selection on 11 rodent branches on which the lifespan was presumably extended – most of them in the clade of the African mole-rat family that covers the longest-lived known rodents (67). The second study examined three branches of the *Nothobranchius* genus on which lifespan was presumably reduced (15) – the genus covers the shortest-lived vertebrate species that can be held in captivity (68).

### Protein homology modeling

Homology modelling of the clownfisch GSTK1 was carried out with SWISS–MODEL (http://swissmodel.expasy.org; (69, 70) using the crystal structures of the dimeric apoform of the human mitochondrial GSTK1 (PDB 3rpp; (56)) and the substrate bound dimer of the rat enzyme PDB 1r4w; (57)). No further optimization was applied to the resulting models. Visualisation, superimposition of the respective crystal structures and the models as well as rendering was carried out using CHIMERA (71).

## Author contributions

AS performed the positive selection analysis and wrote the first draft of the paper, PAP performed the clownfish aclimatation and sampling, MB performed the transcriptome assemblies, MM performed the clownfish survey and helped in writing the paper,, ALS performed the clownfish aclimatation and sampling, JdCR performed the clownfish aclimatation and sampling, MG performed the protein homology modelling, AC concieved and supervised the study, wrote the first draft of the paper. All authors have read and improved the first draft of the paper.

## Acknowledgments

We thank the following individuals at zoos or aquariums for responding to our questionnaire: Vicky Béduneau (Océarium du Croisic, Le Croisic, France), Nicolas Hirel (Aquarium Mare Nostrum, Montpellier, France), Thomas Ziegler (Koelner Zoo, Köln, Germany), Nikolaj Meyer (Skansen-Akvariet, Stockholm, Sweden), Markus Dernjatin (Sea Life Helsinki, Helsinki, Finland). Personal communications on clownfish longevity were received also from Prof. Ike Olivotto (University of Ancona) and Prof. Hellen Thaler (Innsbruck). We thank Cornelia Luge, Ivonne Görlich and Marco Groth (CF DNA sequencing at Leibniz Institute on Aging - Fritz Lipmann Institute) for conducting Illumina sequencing. We thank Matthias Platzer and Steve Hoffmann for helpful discussions and support.

## Supplement

**Supplement Fig. S1.**
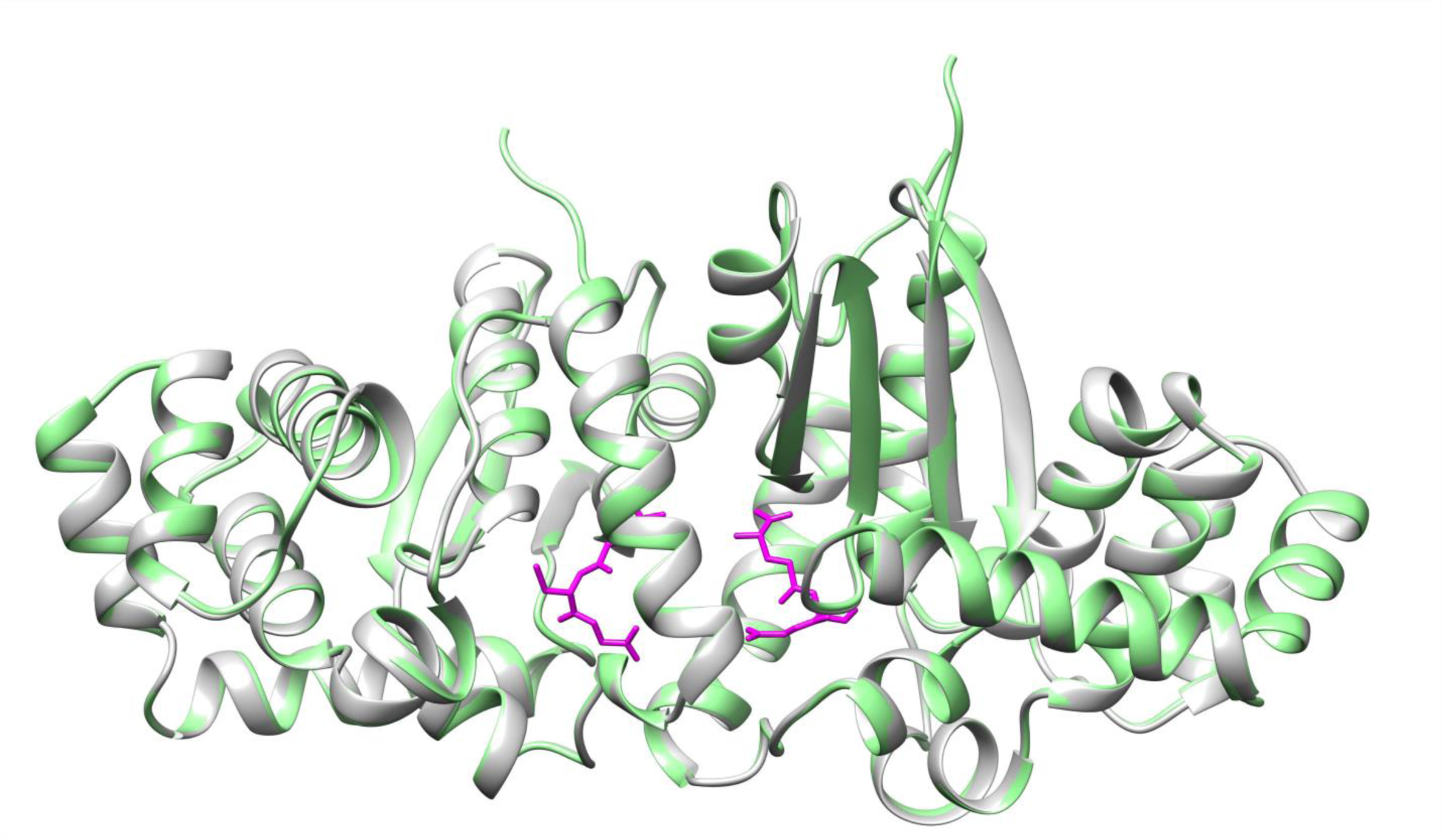
Homology modelling of Clownfish GSTK1. Ribbon representation of the model dimer for the clownfish enzyme as derived from SWISS-MODEL in grey, superimposed onto the dimeric structure of the substrate bound rat GSTK1 (PDB 1r4w; (57)) used as template in light green. The pairwise r.m.s.d. for the Cα positions between the model and 1r4w amounts to 0.52 Å as determined with the CHIMERA Matchmaker tool. The GSH substrate in the rat enzyme structure is rendered in light purple.

### Supplement tables

**Table S1.** Clownfish lifespan questionnaire results.

**Table S2.** PosiGene results for positively selcted genes on the phylogenetic branch representing the last common ancestor of the the clownfishes (genus *Amphiprion*).

**Table S3.** Enrichment test resullts of biological gene ontology processes enriched for positively selected genes.

**Table S4.** Meta analysis using Fisher's method of positive selection across three analyses of phylogenetic branches on which lifespand changed considerably.

**Table S5.** Meta analysis using Fisher's method of positive selection across three phylogenetic branches of the Nothobranchius genus on whichlifespan was reduced considerably.

**Table S6.** Meta analysis using Fisher's method of positive selection across eleven phylogenetic rodent branches on which lifespan was reduced considerably.

**Table S7.** Genes that were regarded as mitochondrial biogenesis related from five gene ontology terms.

**Table S8.** Assembly and sequence statistics.

**Table S9.** Samples that were sequenced to create genome/transcriptome assemblies.

### Supplement data

available at:

ftp://genome.leibniz-fli.de/pub/user/arne.sahm/clownfish/supplement_data.tar.gz> The package contains assembled sequence data, visualizations of alignments and positively selected sites for all genes that were analyzed in this article.

## References

1. Vrtilek M, Zak J, Polacik M, Blazek R, Reichard M. Longitudinal demographic study of wild populations of African annual killifish. Sci Rep. 2018;8(1):4774.

2. Nielsen J, Hedeholm RB, Heinemeier J, Bushnell PG, Christiansen JS, Olsen J, et al. Eye lens radiocarbon reveals centuries of longevity in the Greenland shark (Somniosus microcephalus). Science. 2016;353(6300):702–4.

3. Finch TM, Zhao N, Korkin D, Frederick KH, Eggert LS. Evidence of positive selection in mitochondrial complexes I and V of the African elephant. PLoS One. 2014;9(4):e92587.

4. Seim I, Fang X, Xiong Z, Lobanov AV, Huang Z, Ma S, et al. Genome analysis reveals insights into physiology and longevity of the Brandt’s bat Myotis brandtii. Nat Commun. 2013;4:2212.

5. Keane M, Semeiks J, Webb AE, Li YI, Quesada V, Craig T, et al. Insights into the evolution of longevity from the bowhead whale genome. Cell Rep. 2015;10(1):112–22.

6. Kim EB, Fang X, Fushan AA, Huang Z, Lobanov AV, Han L, et al. Genome sequencing reveals insights into physiology and longevity of the naked mole rat. Nature. 2011;479(7372):223–7.

7. Fang X, Nevo E, Han L, Levanon EY, Zhao J, Avivi A, et al. Genome-wide adaptive complexes to underground stresses in blind mole rats Spalax. Nat Commun. 2014;5:3966.

8. Sahm A, Bens M, Szafranski K, Holtze S, Groth M, Gorlach M, et al. Long-lived rodents reveal signatures of positive selection in genes associated with lifespan. PLoS Genet. 2018;14(3):e1007272.

9. Hossie TJ, Hassall C, Knee W, Sherratt TN. Species with a chemical defence, but not chemical offence, live longer. J Evol Biol. 2013;26(7):1598–602.

10. Blanco MA, Sherman PW. Maximum longevities of chemically protected and non-protected fishes, reptiles, and amphibians support evolutionary hypotheses of aging. Mech Ageing Dev. 2005;126(6-7):794–803.

11. Shattuck MR, Williams SA. Arboreality has allowed for the evolution of increased longevity in mammals. Proc Natl Acad Sci U S A. 2010;107(10):4635–9.

12. Tozzini ET, Dorn A, Ng’oma E, Polacik M, Blazek R, Reichwald K, et al. Parallel evolution of senescence in annual fishes in response to extrinsic mortality. BMC Evol Biol. 2013;13:77.

13. Blazek R, Polacik M, Kacer P, Cellerino A, Rezucha R, Methling C, et al. Repeated intraspecific divergence in life span and aging of African annual fishes along an aridity gradient. Evolution. 2017;71(2):386–402.

14. Cellerino A, Valenzano DR, Reichard M. From the bush to the bench: the annual Nothobranchius fishes as a new model system in biology. Biol Rev Camb Philos Soc. 2016;91(2):511–33.

15. Sahm A, Bens M, Platzer M, Cellerino A. Parallel evolution of genes controlling mitonuclear balance in short-lived annual fishes. Aging Cell. 2017;16(3):488–96.

16. Litsios G, Sims CA, Wuest RO, Pearman PB, Zimmermann NE, Salamin N. Mutualism with sea anemones triggered the adaptive radiation of clownfishes. BMC Evol Biol. 2012;12:212.

17. Fautin DG, Allen GR. Field Guide to Anemonefishes and Their Host Sea Anemones. Perth: Western Australia Museum. 1992.

18. Elliott JK, Mariscal RN. Ontogenetic and interspecific variation in the protection of anemonefishes from sea anemones. J Exp Mar Biol Ecol. 1997;208(1):57–72.

19. Ollerton J, McCollin D, Fautin DG, Allen GR. Finding NEMO: nestedness engendered by mutualistic organization in anemonefish and their hosts. Proc Biol Sci. 2007;274(1609):591–8.

20. Buston PM. Mortality is associated with social rank in the clown anemonefish (Amphiprion percula). Marine Biology 2003;143:811–5.

21. Elliott JK, Elliott JM, Mariscal RN. Host selection, location, and association behaviors of anemonefishes in field settlement experiments. Marine Biology 1995;122(3):377–89.

22. Mariscal RN. The nature of the symbiosis between Indo-Pacific anemone fishes and sea anemones. Mar Biol 1970;6(1):58–65.

23. Buston PM, García MB. An extraordinary life span estimate for the clown anemonefish Amphiprion percula. Journal of Fish Biology. 2007;70(6):1710–9.

24. Aldenhoven JM. Local variation in mortality rates and life-expectancy estimates of the coral reef fish Centropyge bicolor (Pisces: Pomacanthidae). Marine Biology 1986;92:237–44.

25. Eckert GJ. Estimates of adult and juvenile mortality for labrid fishes at One Tree Reef, Great Barrier Reef. Marine Biology 1987;95:161–71.

26. Munro JL, Williams DM. Assessment and management of coral reef fisheries: biological, environmental, and socioeconomic aspects. In Proceedings of the 5th International Coral Reef Congress. 1985;4:545–81.

27. Maison KA, Graham KS. Status review report: orange clownfish (Amphiprion percula). Pacific Islands Fisheries Science Center, National Marine Fisheries Service, National Oceanic and Atmospheric Administration, US Department of Commerce. 2016.

28. Salles OC, Maynard JA, Joannides M, Barbu CM, Saenz-Agudelo P, Almany GR, et al. Coral reef fish populations can persist without immigration. Proc Biol Sci. 2015;282(1819).

29. Sahm A, Platzer M, Cellerino A. Outgroups and Positive Selection: The Nothobranchius furzeri Case. Trends Genet. 2016;32(9):523–5.

30. Garcia-Herrera N, Ferse SCA, Kunzmann A, Genin A. Mutualistic damselfish induce higher photosynthetic rates in their host coral. The Journal of Experimental Biology. 2017;220(10):1803–11.

31. Holbrook SJ, Brooks AJ, Schmitt RJ, Stewart HL. Effects of sheltering fish on growth of their host corals. Mar Biol. 2008;155(5):521–30.

32. Sweatman H. The timing of settlement by larval Dascyllus aruanus: some consequences for larval habitat selection. Proc 5th Int Coral Reef Conf 5. 1985:367–72.

33. Liberman T, Genin A, Loya Y. Effects on growth and reproduction of the coral Stylophora pistillata by the mutualistic damselfish Dascyllus marginatus. Mar Biol. 1995;121:741–6.

34. Ben-Tzv iO, Abelson A, Polak O, Kiflawi M. Habitat selection and the colonization of new territories by Chromis viridis. Journal of Fish Biology. 2008;73(4):1005–18.

35. Lecchini D, Nakamura Y, Grignon J, Tsuchiya M. Evidence of density-independent mortality in a settling coral reef damselfish, Chromis viridis. Ichthyological Research. 2006;53(3):298–300.

36. Hixon MA, Carr MH. Synergistic Predation, Density Dependence, and Population Regulation in Marine Fish. Science. 1997;277(5328):946–9.

37. Moyer JT. Longevity of the Anemonefish Amphiprion clarkii at Miyake-Jima, Japan with Notes on Four Other Species Copeia. 1986;1986:135–9.

38. McElwee JJ, Schuster E, Blanc E, Piper MD, Thomas JH, Patel DS, et al. Evolutionary conservation of regulated longevity assurance mechanisms. Genome Biol. 2007;8(7):R132.

39. Swindell WR. Gene expression profiling of long-lived dwarf mice: longevity-associated genes and relationships with diet, gender and aging. BMC Genomics. 2007;8:353.

40. Amador-Noguez D, Dean A, Huang W, Setchell K, Moore D, Darlington G. Alterations in xenobiotic metabolism in the long-lived Little mice. Aging Cell. 2007;6(4):453–70.

41. Steinbaugh MJ, Sun LY, Bartke A, Miller RA. Activation of genes involved in xenobiotic metabolism is a shared signature of mouse models with extended lifespan. Am J Physiol Endocrinol Metab. 2012;303(4):E488–95.

42. Houtkooper RH, Mouchiroud L, Ryu D, Moullan N, Katsyuba E, Knott G, et al. Mitonuclear protein imbalance as a conserved longevity mechanism. Nature. 2013;497(7450):451–7.

43. Valenzano DR, Benayoun BA, Singh PP, Zhang E, Etter PD, Hu CK, et al. The African Turquoise Killifish Genome Provides Insights into Evolution and Genetic Architecture of Lifespan. Cell. 2015;163(6):1539–54.

44. Sahm A, Cellerino A. (Anti-)parallel evolution of lifespan. Aging (Albany NY). 2017;9(10):2018–9.

45. Antonicka H, Shoubridge EA. Mitochondrial RNA Granules Are Centers for Posttranscriptional RNA Processing and Ribosome Biogenesis. Cell Rep. 2015.

46. Eskelinen EL. Roles of LAMP-1 and LAMP-2 in lysosome biogenesis and autophagy. Mol Aspects Med. 2006;27(5-6):495–502.

47. Berditchevski F, Odintsova E. Tetraspanins as regulators of protein trafficking. Traffic. 2007;8(2):89–96.

48. Sordat I, Decraene C, Silvestre T, Petermann O, Auffray C, Pietu G, et al. Complementary DNA arrays identify CD63 tetraspanin and alpha3 integrin chain as differentially expressed in low and high metastatic human colon carcinoma cells. Lab Invest. 2002;82(12):1715–24.

49. Sauer G, Kurzeder C, Grundmann R, Kreienberg R, Zeillinger R, Deissler H. Expression of tetraspanin adaptor proteins below defined threshold values is associated with in vitro invasiveness of mammary carcinoma cells. Oncol Rep. 2003;10(2):405–10.

50. Zhijun X, Shulan Z, Zhuo Z. Expression and significance of the protein and mRNA of metastasis suppressor gene ME491/CD63 and integrin alpha5 in ovarian cancer tissues. Eur J Gynaecol Oncol. 2007;28(3):179–83.

51. Kwon MS, Shin SH, Yim SH, Lee KY, Kang HM, Kim TM, et al. CD63 as a biomarker for predicting the clinical outcomes in adenocarcinoma of lung. Lung Cancer. 2007;57(1):46–53.

52. Lai X, Gu Q, Zhou X, Feng W, Lin X, He Y, et al. Decreased expression of CD63 tetraspanin protein predicts elevated malignant potential in human esophageal cancer. Oncol Lett. 2017;13(6):4245–51.

53. Zhang C, Cuervo AM. Restoration of chaperone-mediated autophagy in aging liver improves cellular maintenance and hepatic function. Nat Med. 2008;14(9):959–65.

54. Gao F, Fang Q, Zhang R, Lu J, Lu H, Wang C, et al. Polymorphism of DsbA-L gene associates with insulin secretion and body fat distribution in Chinese population. Endocr J. 2009;56(3):487–94.

55. Sharma M, Gupta S, Singh K, Mehndiratta M, Gautam A, Kalra OP, et al. Association of glutathione-S-transferase with patients of type 2 diabetes mellitus with and without nephropathy. Diabetes Metab Syndr. 2016;10(4):194–7.

56. Wang B, Peng Y, Zhang T, Ding J. Crystal structures and kinetic studies of human Kappa class glutathione transferase provide insights into the catalytic mechanism. Biochem J. 2011;439(2):215–25.

57. Ladner JE, Parsons JF, Rife CL, Gilliland GL, Armstrong RN. Parallel evolutionary pathways for glutathione transferases: structure and mechanism of the mitochondrial class kappa enzyme rGSTK1-1. Biochemistry. 2004;43(2):352–61.

58. Aramillo Irizar P, Schauble S, Esser D, Groth M, Frahm C, Priebe S, et al. Transcriptomic alterations during ageing reflect the shift from cancer to degenerative diseases in the elderly. Nat Commun. 2018;9(1):327.

59. Kurz T, Terman A, Gustafsson B, Brunk UT. Lysosomes and oxidative stress in aging and apoptosis. Biochim Biophys Acta. 2008;1780(11):1291–303.

60. Bens M, Sahm A, Groth M, Jahn N, Morhart M, Holtze S, et al. FRAMA: from RNA-seq data to annotated mRNA assemblies. BMC Genomics. 2016;17:54.

61. Simao FA, Waterhouse RM, Ioannidis P, Kriventseva EV, Zdobnov EM. BUSCO: assessing genome assembly and annotation completeness with single-copy orthologs. Bioinformatics. 2015;31(19):3210–2.

62. Sahm A, Bens M, Platzer M, Szafranski K. PosiGene: automated and easy-to-use pipeline for genome-wide detection of positively selected genes. Nucleic Acids Res. 2017.

63. Overbeek R, Fonstein M, D’Souza M, Pusch GD, Maltsev N. The use of gene clusters to infer functional coupling. Proc Natl Acad Sci U S A. 1999;96(6):2896–901.

64. Camacho C, Coulouris G, Avagyan V, Ma N, Papadopoulos J, Bealer K, et al. BLAST+: architecture and applications. BMC Bioinformatics. 2009;10:421.

65. Benjamini Y, Y. H. Controlling the false discovery rate: A practical and powerful approach to multiple testing. Journal of the Royal Statistical Society Series B (Methodological). 1995;57(1):289–300.

66. Fisher RA. Statistical Methods for Research Workers. 1932.

67. Tacutu R, Craig T, Budovsky A, Wuttke D, Lehmann G, Taranukha D, et al. Human Ageing Genomic Resources: integrated databases and tools for the biology and genetics of ageing. Nucleic Acids Res. 2013;41(Database issue):D1027–33.

68. Valdesalici S, Cellerino A. Extremely short lifespan in the annual fish Nothobranchius furzeri. Proc Biol Sci. 2003;270 Suppl 2:S189–91.

69. Arnold K, Bordoli L, Kopp J, Schwede T. The SWISS-MODEL workspace: a web-based environment for protein structure homology modelling. Bioinformatics. 2006;22(2):195–201.

70. Biasini M, Bienert S, Waterhouse A, Arnold K, Studer G, Schmidt T, et al. SWISS-MODEL: modelling protein tertiary and quaternary structure using evolutionary information. Nucleic Acids Res. 2014;42(Web Server issue):W252–8.

71. Pettersen EF, Goddard TD, Huang CC, Couch GS, Greenblatt DM, Meng EC, et al. UCSF Chimera−- a visualization system for exploratory research and analysis. J Comput Chem. 2004;25(13):1605–12.

72. Felsenstein J. PHYLIP (Phylogeny Inference Package) version 3.6. Distributed by the author. 2005; Department of Genome Sciences, University of Washington, Seattle.

